# Shared Spatiotemporal Category Representations in Biological and Artificial Deep Neural Networks

**DOI:** 10.1101/225607

**Authors:** Michelle R. Greene, Bruce C. Hansen

## Abstract

Understanding the computational transformations that enable invariant visual categorization is a fundamental challenge in both systems and cognitive neuroscience. Recently developed deep convolutional neural networks (CNNs) perform visual categorization at accuracies that rival humans, providing neuroscientists with the opportunity to interrogate the series of representational transformations that enable categorization *in silico*. The goal of the current study is to assess the extent to which sequential visual representations built by a CNN map onto those built in the human brain as assessed by high-density, time-resolved event-related potentials (ERPs). We found correspondence both over time and across the scalp: earlier ERP activity was best explained by early CNN layers at all electrodes. Later neural activity was best explained by the later, conceptual layers of the CNN. This effect was especially true both in frontal and right occipital sites. Together, we conclude that deep artificial neural networks trained to perform scene categorization traverse similar representational stages as the human brain. Thus, examining these networks will allow neuroscientists to better understand the transformations that enable invariant visual categorization.

## Introduction

Categorization, the act of grouping like with like, is a hallmark of human intelligence. Visual recognition of scene environments in particular is incredibly rapid (Greene & Oliva, 2009; Potter, Wyble, Hagmann, & McCourt, 2014), and may even be automatic (Greene & Fei-Fei, 2014). Despite the importance of this problem, little is known about the temporal dynamics of neural activity that give rise to categorization. A common conceptual framework in the visual neuroscience community considers representations in each visual area as a geometric space, with individual images represented as points within this representational space (Edelman, 1998; Kriegeskorte, Mur, & Bandettini, 2008). According to this account, early visual processing can be described as a tangled geometric surface in which individual categories cannot be easily separated, and that over the course of processing, the ventral visual stream disentangles these manifolds allowing for categories to be distinguished (DiCarlo & Cox, 2007; Riesenhuber & Poggio, 1999). Although this view has provided a useful descriptive framework, we still know little about how this disentangling occurs because it has been difficult to examine the representations at each stage of processing.

In a parallel development, work in computer vision has resulted in the creation of deep convolutional neural networks (CNNs) whose categorization abilities rival those of human observers (Russakovsky et al., 2015). Although CNNs do not explicitly seek to model the human visual system, their architectures are inspired by the structure of the human visual system (Fukushima, 1988): like the human visual system, they are arranged hierarchically in discrete representational layers, and they apply both linear and nonlinear operations across the whole of visual space. As CNNs are explicitly trained for the purpose of visual categorization, and because they achieve near human-level performance at this task, they provide neuroscientists with an unprecedented opportunity to interrogate the types of intermediate level representations that are built en route to categorization. Indeed, there is a growing literature detailing the similarities between aspects of CNN activity and activity in biological brains (Agrawal, Stansbury, Malik, & Gallant, 2014; Cadieu et al., 2014; Cichy, Khosla, Pantazis, Torralba, & Oliva, 2016; Güçlü & Gerven, 2015; Khaligh-Razavi & Kriegeskorte, 2014; Kubilius, Bracci, & Beeck, 2016; Yamins, Hong, Cadieu, & DiCarlo, 2013; Yamins & DiCarlo, 2016).

Of particular interest to the current study are the correspondences between the stages of processing within a CNN and the temporal order of processing in the human brain, as assessed with M/EEG. Although studies have demonstrated that the brain activity elicited by individual images can be predicted by the CNN (Cichy et al., 2016; Seeliger et al., 2017), and that upper layers of the CNN can predict semantically-relevant spatial properties such as overall scene volume (Cichy, Khosla, Pantazis, & Oliva, 2017), the issue of scene category membership, and how intermediate stages of visual representation allow for these complex abstractions to take place, remains open. Furthermore, these studies pool across all sensors at a given time point, discarding potentially informative scalp patterns. While this capitalizes on the fine temporal scale of M/EEG, a complete understanding of the neural dynamics of scene understanding requires the characterization of information flow across the cortex.

Therefore, the goal of the current study is to assess the extent to which sequential representations in each layer of a pre-trained deep convolutional neural network predict the sequential representations built by the human brain using high-density, time-resolved event-related potentials (ERPs). Previewing our results, we show a spatiotemporal correspondence between sequential CNN layers and the order of processing in the human visual system. Specifically, earlier layers of the CNN correspond better to early time points in human visual processing, and to electrodes over occipital and left temporal scalp regions. Later layers of the CNN correspond to later information, and best match electrodes in the frontal half of the scalp, and in right occipitotemporal cortex. The correspondence between computer models and human vision provides neuroscientists with the unique opportunity to probe intermediate-level representations *in silico*, allowing for a more complete understanding of the neural computations underlying visual categorization.

## Methods

### Participants

Fourteen observers (6 female, 13 right-handed) participated in the study. One of the participant’s data contained fewer than half valid trials following artifact rejection and was removed from all analyses. An additional 15 participants (9 female, 12 right-handed) were recruited as part of an internal replication study (results from those participants are reported as Supplementary Material). The age of all participants ranged from 18 to 22 (mean age = 19.1 for main experiment, and 19.4 for replication study). All participants had normal (or corrected to normal) vision as determined by standard ETCRS acuity charts. All participants gave Institutional Review Board-approved written informed consent before participating, and were compensated for their time.

### Stimuli

The stimuli consisted of 2250 color images of real-world photographs from 30 different scene categories (75 exemplars in each category) taken from the SUN database (Xiao, Ehinger, Hays, Torralba, & Oliva, 2014). When possible, images were taken from the SUN database. In cases where this database did not have 75 images, we sampled from the internet (copyright-free images). Care was taken to omit images with salient faces in them. All images had a resolution of 512 x 512 pixels (which subtended 20.8° of visual angle) and were processed to possess the same root-mean-square (RMS) contrast (luminance and color) as well as mean luminance. All images were fit with a circular linear edge-ramped window to obscure the square frame of the images, thereby uniformly distributing contrast changes around the circular edge of the stimulus (Hansen & Essock, 2004; Hansen & Hess, 2006).

### Apparatus

All stimuli were presented on a 23.6” VIEWPixx/EEG scanning LED-backlight LCD monitor with 1ms black-to-white pixel response time. Maximum luminance output of the display was 100 cd/m^2^, with a frame rate of 120 Hz and resolution of 1920 x 1080 pixels. Single pixels subtended 0.0406° of visual angle (i.e. 2.43 arc min.) as viewed from 32 cm. Head position was maintained with an Applied Science Laboratories (ASL) chin rest.

### Experimental Procedure

Participants engaged in a 3 alternative forced-choice (3AFC) categorization task with each of the 2250 images. As it was not feasible for participants to view all images in one sitting, all images were randomly split into two sets, keeping equal numbers of images within each category. Each image set was presented within a different ~50-minute recording session, run on separate days. The image set was counterbalanced across participants, and image order within each set was randomized. Participants viewed the windowed scenes against a mean luminance background under darkened conditions (i.e. the only light source in the testing chamber was the monitor). All trials began with a 500 ms fixation followed by a variable duration (500-750 ms) blank mean luminance screen to enable any fixation-driven activity to dissipate. Next, a scene image was presented for 750 ms followed by a variable 100-250 ms blank mean luminance screen, followed by a response screen consisting of the image’s category name and the names of two distractor categories presented laterally in random order (distractor category labels were randomly sampled from the set of 29 and therefore varied on a trial-by-trial basis). Observers selected their choice via mouse click. Performance feedback was not given.

### EEG Recording and Processing

Continuous EEGs were recorded in a Faraday chamber using EGI’s Geodesic EEG acquisition system (GES 400) with Geodesic Hydrocel sensor nets consisting of a dense array of 256 channels (electrolytic sponges). The on-line reference was at the vertex (Cz), and the impedances were maintained below 50 kΩ (EGI amplifiers are high-impedance amplifiers – this value is optimized for this system). All EEG signals were amplified and sampled at 1000 Hz. The digitized EEG waveforms were band-pass filtered offline from 0.1 Hz to 45 Hz to remove the DC offset and eliminate 60 Hz line noise.

All continuous EEGs were divided into 850 msec epochs (100 ms before stimulus onset and the 750 ms of stimulus-driven data). Trials that contained eye movements or eye blinks during data epochs were excluded from analysis. Further, all epochs were subjected to algorithmic artifact rejection of voltage exceeding ± 100 μV. These trial rejection routines resulted in no more than 10% of the trials being rejected on a participant-by-participant basis. Each epoch was then rereferenced offline to the net average, and baseline corrected to the last 100 msec of the luminance blank interval that preceded the image. Grand average event-related potentials (ERPs) were assembled by averaging all re-referenced and baseline corrected epochs across participants. Topographic plots were generated for all experimental conditions using EEGLAB (Delorme & Makeig, 2004) version 13.4.4b in MATLAB (ver. R2016a, The Mathworks, MA).

For all analyses, we improved the signal-to-noise ratio of the single trial data by using a bootstrapping approach to build sub-averages across trials for each trial (e.g., Cichy, Khosla, Pantazis, & Oliva, 2016). Specifically, for each trial within a given scene category, we randomly selected 20% of the trials within that category and averaged those to yield a sub-averaged ERP for that trial. This was repeated until all valid trials within each category were built. This process was repeated separately for each participant. This approach is desirable as we are primarily interested in category-level neuroelectric signals that are time-locked to the stimulus.

### Deep Convolutional Neural Network (CNN)

In order to assess the representations available in a deep convolutional neural network at each processing stage, we extracted the activations in each of eight layers in a pre-trained network. Specifically, we used a CNN based on the AlexNet architecture (Krizhevsky, Sutskever, & Hinton, 2012) that was pre-trained on the Places database (Zhou, Lapedriza, Khosla, Oliva, & Torralba, 2017) and implemented in Caffe (Jia et al., 2014). The first five layers of this neural network are convolutional, and the last three are fully connected. This CNN was chosen because it is optimized for 205-category scene classification, and because the eight-layer architecture, loosely inspired by biological principles, is most frequently used when comparing CNNs and brain activity (Cichy et al., 2016; Kubilius et al., 2016; Yamins et al., 2013). For each layer, we averaged across images within a category, creating 30-category by N-feature matrices.

In order to assess the amount of category-related information available in each layer of the CNN, we performed a decoding procedure on the feature vectors from each CNN layer using a linear multi-class support vector machine (SVM) implemented as LIBSVM in Matlab (Chang & Lin, 2011). Decoding accuracies for each layer were calculated using 5-fold cross validation.

For all analyses, statistical testing was done via permutation testing. Specifically, we randomly shuffled the RDMs (1000 permutation samples per participant) to create an empirical chance distribution. To correct for multiple comparisons, we used cluster extent with a threshold of p<0.05, Bonferroni-corrected for multiple comparisons (similar to Cichy et al., 2016).

### Time-Resolved Encoding Analysis

Our approach is graphically illustrated in **Figure 1.** To compare category representations in the CNN to those of human observers, we used the representational similarity analysis (RSA, Kriegeskorte et al., 2008). This allowed us to directly compare models and neuroelectric signals by abstracting both into a similarity space.

**Figure 1:**
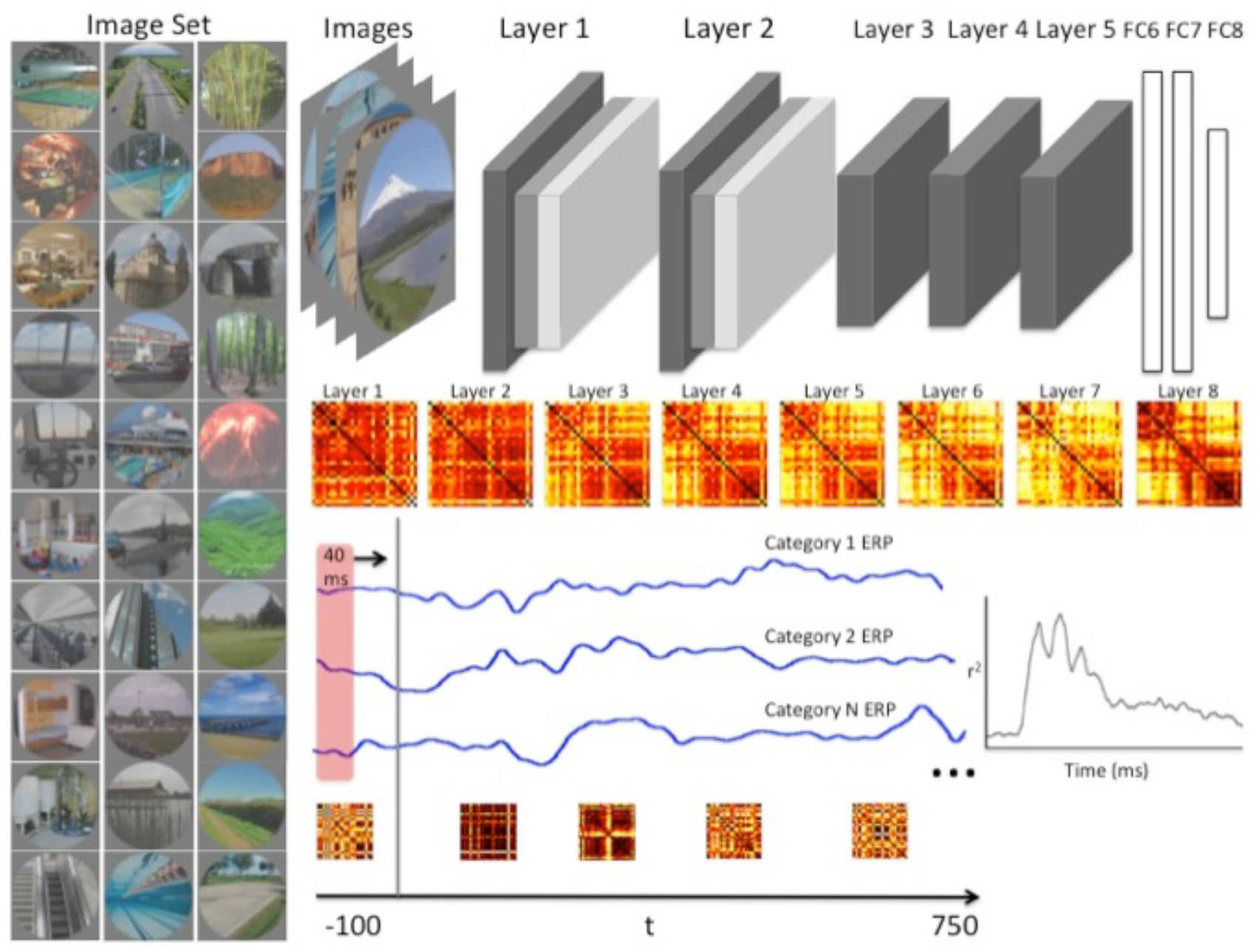
*The image set consisted of 75 images from each of 30 scene categories. Example images are shown on the left. Activations for each of the eight layers of a pre-trained deep convolutional neural network (CNN) were extracted for each of the 2250 images. We averaged across exemplars to create 30x30 representational dissimilarity matrices (RDMs) for each layer of the CNN. For each participant, and for each of the 256 electrodes, we ran a 40 ms sliding window from the 100 ms before stimulus presentation, and running through the 750 ms where image was on screen. At each time point, a 30 x 30 RDM was created and compared to each layer of the CNN using correlation. Note: although full matrices are shown, only the lower triangle was used in analysis*.

For each of the eight layers of the CNN, we created 30-category by 30-category correlation matrices. Representational dissimilarity matrices (RDMs) were created by transforming the correlation matrices into distance matrices using the metric of 1-Spearman rho (Cichy et al., 2016; Khaligh-Razavi & Kriegeskorte, 2014). Each RDM is a symmetrical matrix with an undefined diagonal. Therefore, in all subsequent analyses, we will use only the lower triangle of each RDM to represent patterns of category similarity.

To create neural RDMs, for each participant and for each electrode, we extracted ERP signals within a 40 ms sliding window beginning 100 ms before image presentation, and extending to the entire 750 ms image duration. For each window, we created a 30 x 30 RDM using the same 1-minus-correlation distance metric described above. The window size was truncated at the end of each trial as to not extend beyond image presentation. The upper and lower bounds of the noise ceiling for the data were computed as recommended in (Nili et al., 2014). It is worth noting that while previous MEG RDM results have performed time-resolved analyses on a time point by time point manner taking the value at each sensor at each time point as a feature (e.g. Cichy et al., 2016), our approach allows for the understanding of feature correspondence at each electrode, also enabling spatiotemporal analysis.

## Electrode Clustering

In order to examine spatial relations among encoding patterns across electrodes, we adopted a data-driven approach based on random field theory. A similar approach has been used to assess statistical significance of pixel clusters in classification images (Chauvin, Worsley, Schyns, Arguin, & Gosselin, 2005). This allowed us to identify spatially contiguous electrode clusters based on voltage differences, while remaining agnostic to any encoding differences with the CNN. Specifically, we submitted participant-averaged and z-transformed voltage difference topographic maps to the algorithm time point by time point. The spatial clustering algorithm then grouped electrodes that contained normalized voltage differences that significantly deviated from baseline noise (p < .001). As this analysis was run time point by time point, we selected clusters that persisted for more than 20 ms, resulting in five clusters: an early central occipital cluster (100-125 ms), an early central cluster (80-105 ms), two bilateral occiptotemporal clusters (70-500 ms), and a large frontal cluster (135-500 ms). In subsequent analyses, we will assess the relative encoding strength of each of the eight CNN layers within each of these five clusters.

## Results

### CNN Decoding Results

In order to understand the potential contributions of each CNN layer to category-specific neural responses, we assessed the extent to which each layer contained decodable category information. All CNN layers contained above-chance information for decoding the 30 scene categories (see **Figure 2**). The classification accuracy ranged from 40.8% in the first layer (Conv1) to 90.3% correct in the first fully connected layer (FC6). Somewhat unexpectedly, this is higher classification than that of the top layer (FC8: 84.6%). It is also noteworthy that this level of classification is higher than the 69-70% top-1 classification accuracy reported in (Zhou et al., 2017). This is likely due to the fact that some of the images in our dataset were from the training set of this CNN. In sum, we can expect to see category-specific information in each of the eight layers of the CNN, with maximum category information coming from the fully connected layers.

**Figure 2:**
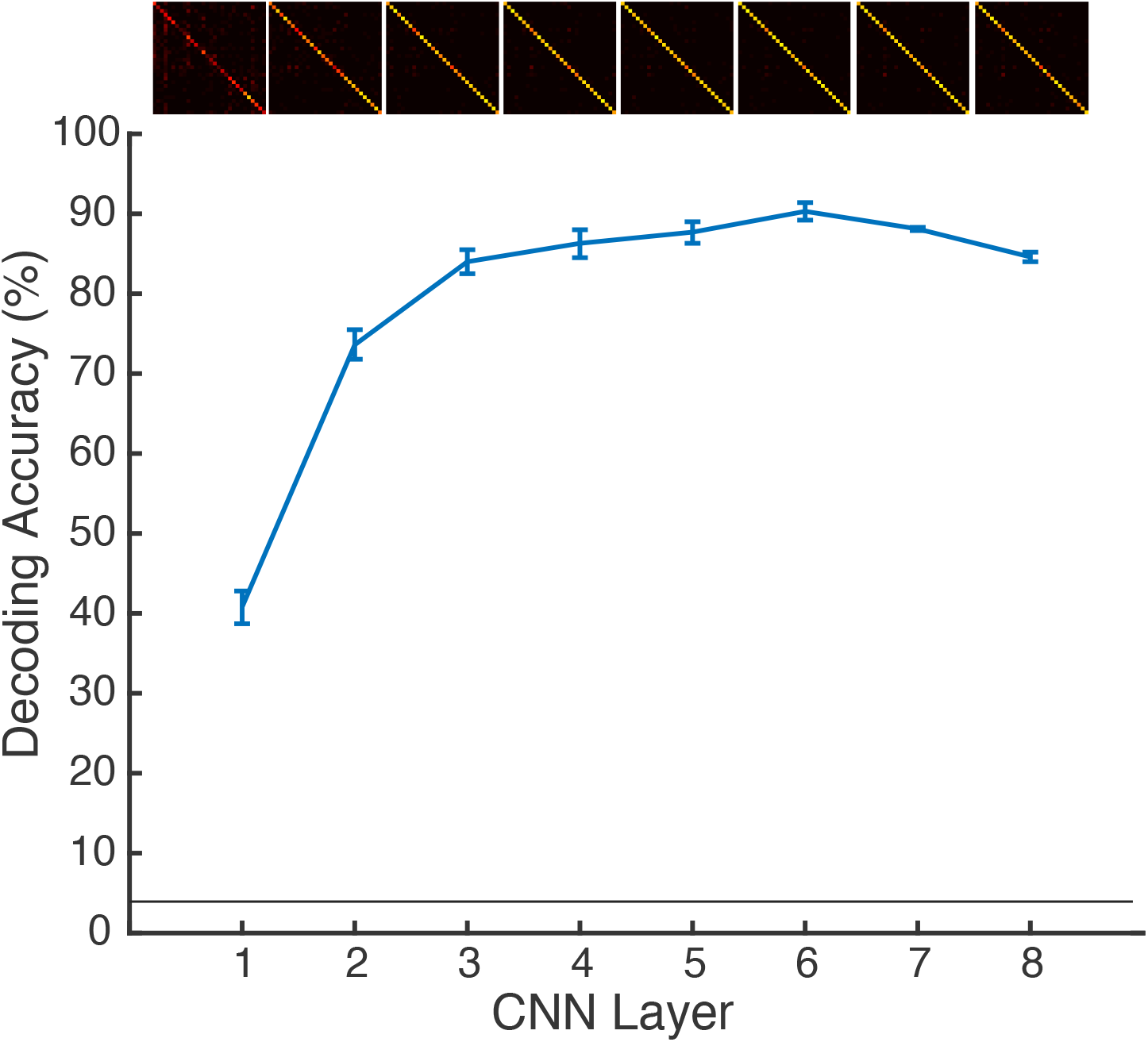
*Category decoding accuracy for each layer of the CNN along with decoding confusion matrices (top row). Error bars reflect 95% confidence intervals. Horizontal line indicates chance level (1/30)*.

### Time-resolved Encoding Results

To get an overall picture of human-CNN correspondence, we examined the extent to which the set of all CNN layers have explanatory power for visual ERPs. Our data-driven clustering method identified five groups of electrodes. We averaged ERPs within each cluster, and then examined the variance explained by all layers of the CNN within that cluster’s RDMs over time. For each cluster, we examined the onset of significant explained variance, the maximum explained variance, and the latency of peak explained variance. We observed no statistically significant differences between the clusters for onset (F(4,60)<1), peak explained variance (F(4,60)=1.11, p=0.36, nor latency at peak (F(4,60)<1). Thus, we show the average of all 256 electrodes in **Figure 3**. We found that the CNN could predict activity starting at 54 ms after scene presentation, and achieved maximal predictive power 93 ms after stimulus onset.

**Figure 3:**
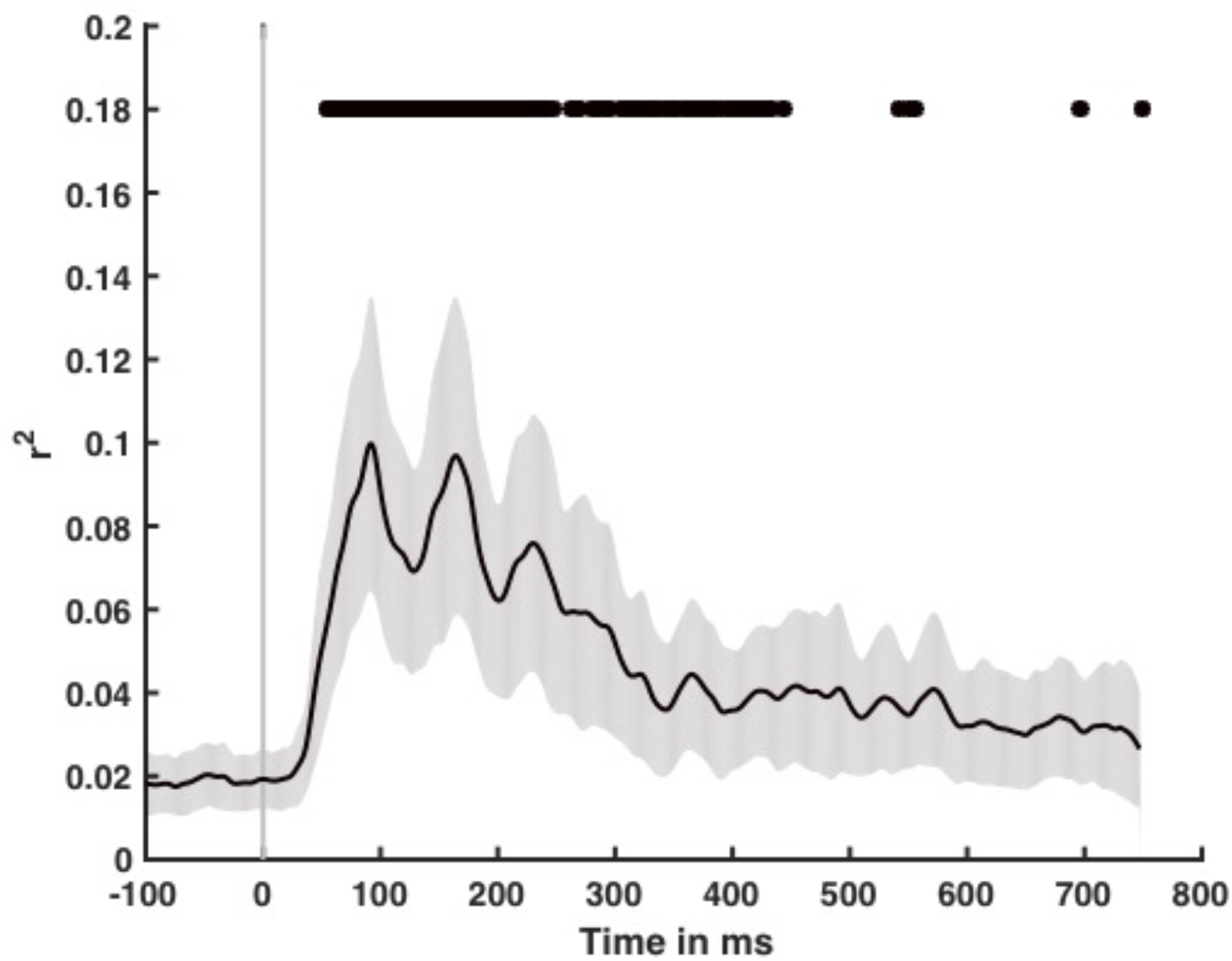
*Variance explained over time for all eight CNN layers together for the average of all 256 electrodes. Black line indicates mean with shaded gray region reflecting 95% confidence interval. Top line indicates statistical significance*.

How much of the explainable ERP variance is the CNN capturing? We estimated the noise ceiling for these ERP data using a method suggested by (Nili et al., 2014), and found that the maximum explainable variance to be 48-54%. Therefore, the maximum *r*^2^of 0.10 across all electrodes does not account for all variability in this dataset. As CNNs are exclusively feedforward models, this result emphasizes the role of feedback in human scene categorization (Bar et al., 2006; Clarke, Devereux, Randall, & Tyler, 2015).

In order to examine the correspondence between the processing stages of the CNN and the representational stages in human visual processing, we next examined the variance explained by each individual CNN layer. Averaged across all 256 electrodes, we observed a correspondence between the onset of explained variability and CNN layer, with earlier layers explaining ERP variability earlier than later layers (r=0.38, p<0.001, see **Figure 4**). Additionally, we observed a negative relationship between CNN layer and the amount of explained variability, with earlier layers explaining more ERP variability than later layers. This effect was pronounced in the first 100 ms post-stimulus (r=-0.51, p<0.0001), and was also observed between 101 and 200 ms (r=-0.26, p<0.005).

**Figure 4:**
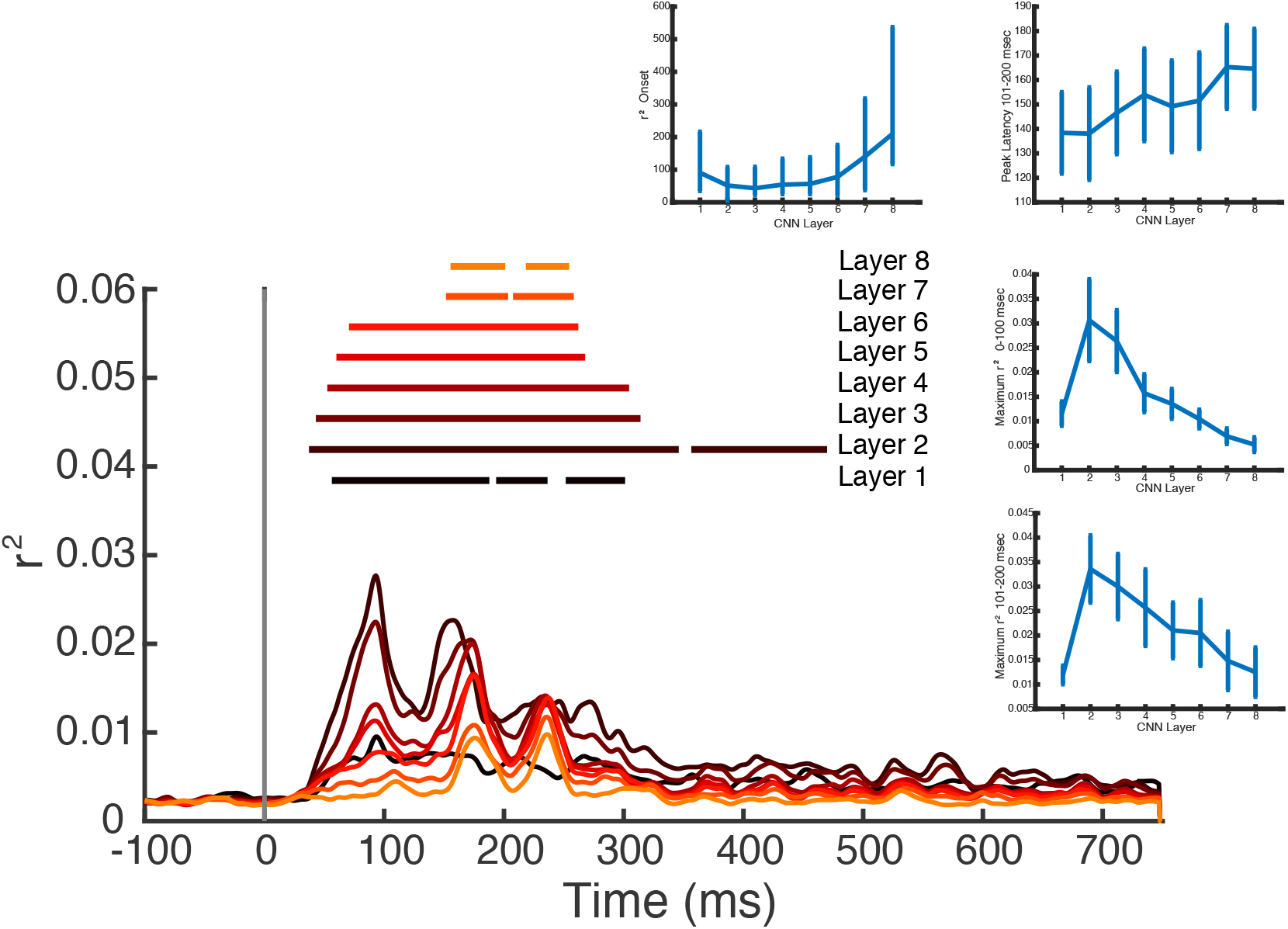
*Main plot: variance explained in each of the eight layers of the CNN taken alone. Each waveform represents the average of all 256 electrodes. Lowest layers are in darker colors, and horizontal lines represent statistical significance. Insets, clockwise from top: Onset of statistically significant encoding as a function of CNN layer; peak variance explained between 101 and 200 ms as a function of CNN layer; maximum variance explained between 0 and 100 ms as a function of CNN layer; maximum variance explained between 101 and 200 ms as a function of CNN layer. Taken together, we can see that early CNN layers explain more variance in ERP patterns, and do so earlier than later CNN layers*.

A chief advantage of conducting the encoding analyses at each electrode, rather than collapsing across all sensors as has been done in previous work (Cichy et al., 2016; Seeliger et al., 2017) is that we are able to examine the spatiotemporal patterns of explained variability, rather than just temporal. Using data-driven electrode clustering, we identified five groups of spatially contiguous electrodes with differing voltage patterns. In each group, we examined the maximum explained variability for each layer of the CNN. As shown in **Figure 5**, we observed a striking dissociation. While the central occipital and left occipitotemporal clusters were better explained by lower layers of the CNN at all time points, the right occipitotemporal cluster was best explained by early layers in early time bins (80-110 ms), by mid-level layers between 120-200 ms, and by the later layers between 200-250 ms post stimulus onset. Similarly, while the frontal cluster best reflected lower layers of the CNN early, it best reflected information from the sixth CNN layer (FC6) in the 200-250 ms time bin. Interestingly, this layer had the maximum decodable category information (see **Figure 2**), suggesting that these signals reflect category-specific information.

**Figure 5:**
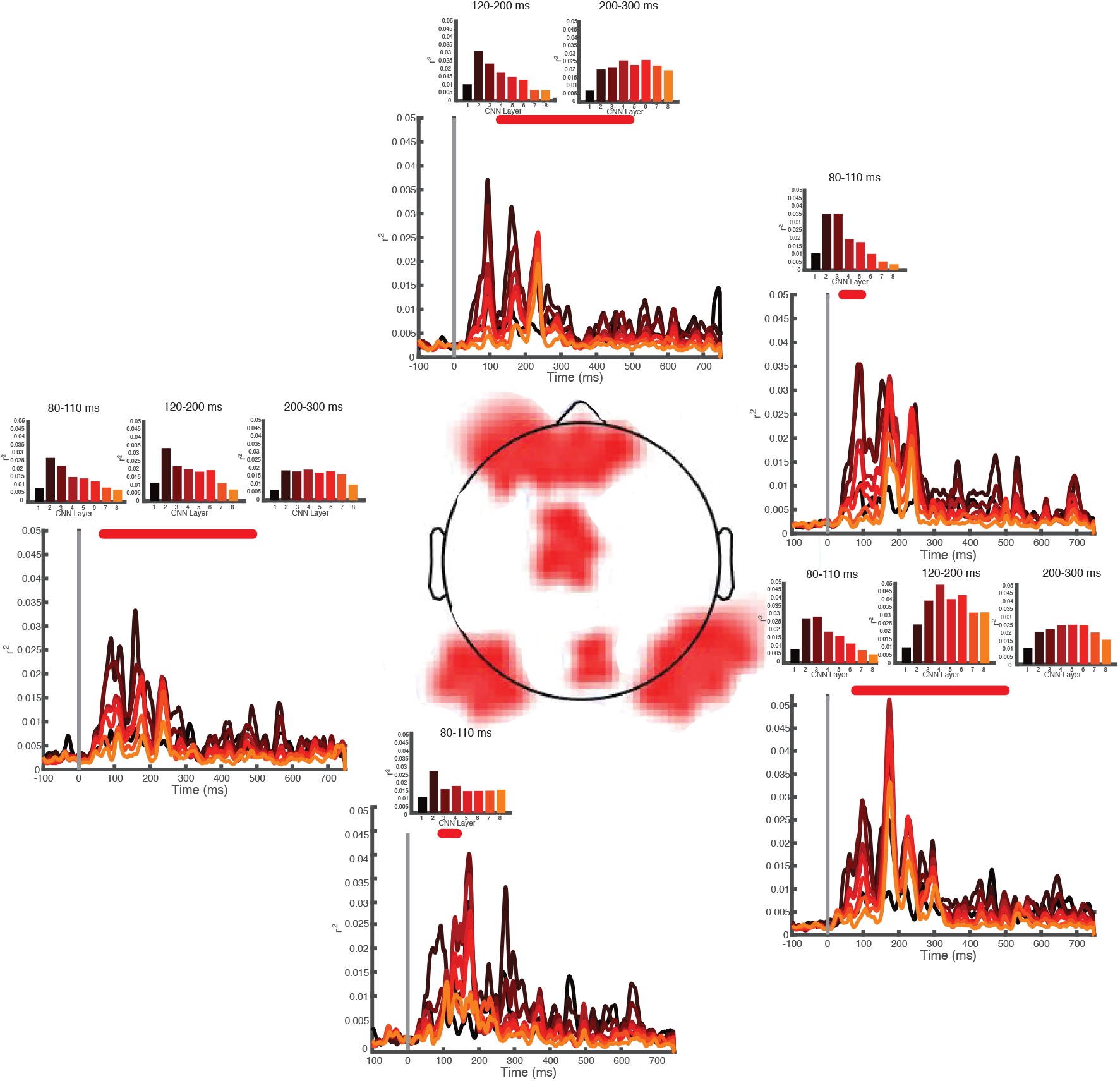
*Encoding time course for each of the five clusters identified. Red lines indicate the times of electrode cluster cohesion, and bar plots indicate maximum explained variability within individual peaks within the cohesive windows*.

## Discussion

In this work, we demonstrated that there is a substantial resemblance between the sequential representations from a pre-trained deep convolutional neural network (CNN), and those in human ERP activity while observers are engaged in categorization. Early layers of the CNN best predicted early ERP activity, while later layers best predicted later activity. Furthermore, our results speak to the importance of feedback processing in human scene categorization as the total variability captured by the feedforward CNN was slightly less than half of the variability given by the noise ceiling of the data, and earlier CNN layers were more predictive of neural activity than the later layers.

Furthermore, we observed a spatial correspondence between CNN layers and ERP variability. While electrodes over central- and left-occipitotemporal cortex were robustly predicted early by early CNN layers, electrodes over right occipitotemporal cortex had sequential representations that resembled the sequential representations of the CNN. Specifically, activity in these electrodes was best predicted by the second and third CNN layers before 100 ms post-stimulus, by the fourth layer between 120-200 ms, and by the sixth layer after 200 ms. A similar striking dissociation was observed over the frontal electrode cluster: early CNN layers best predicted early activity, but the more conceptual, fully-connected layers captured activity at the frontal electrodes about 100 ms later. Taken together, these results suggest a deeper homology between the human visual system and the deep neural network than the few biological principles that inspired the architecture (Krizhevsky et al., 2012). This homology provides the unique intellectual opportunity to probe the mid- and late-stages of visual representation that have thus far been difficult to ascertain.

Because the CNN is an exclusively feedforward model, comparing the representations in the CNN and the human brain afforded us the opportunity to probe the extent to which human category representations are processed in a feedforward manner. Toward that end, we observed that the neural variability explained by the CNN only reached about half of the maximum possible explained variability given by the noise ceiling. This suggests that feedback or recurrent processing plays a significant role in categorization (Bar et al., 2006; Goddard, Carlson, Dermody, & Woolgar, 2016; Hochstein & Ahissar, 2002). Furthermore, we observed that earlier layers of the CNN explained more ERP variance compared with later layers of the CNN. This may be evidence that earlier visual processing is predominantly feedforward, while later visual processing requires feedback.

Our results are largely in agreement with previous MEG studies that examined the correspondence between CNNs and time-resolved neural signals (Cichy et al., 2016; Seeliger et al., 2017). These studies examined whole-brain signals in a time point by time point manner, losing any spatial response pattern. By contrast, by using a sliding window on each electrode, we were able to retain the patterns across the scalp while still examining time-resolved data. Given the known representational differences between MEG and EEG (Cichy & Pantazis, 2017), this work provides unique but complementary information about visual category representations. Specifically, Cichy and Pantazis (2017) found that compared to EEG, MEG signals had decodable information earlier, and more driven by early visual cortex. This suggests that the time points reported here might constitute an upper bound on information availability.

A second difference between this work and previous is that we examined category-specific information instead of image-level information. As the act of recognition is generally an act of categorization (Bruner, 1957), and because the CNN we used was pre-trained specifically to make category-level distinctions (LeCun, Bengio, & Hinton, 2015; Zhou, Lapedriza, Xiao, Torralba, & Oliva, 2014), we argue that this is the most natural comparison for human and artificial neural networks. Accordingly, we assessed the amount of category-level information available in each layer of the CNN. While it is unsurprising that significant category information exists in all eight layers, or that the amount of information increases across layers, we were surprised to observe that category information peaked in the sixth layer. Given that the utility of layer depth is still controversial within the computer vision community (Ba & Caruana, 2014; He, Zhang, Ren, & Sun, 2016), this result may be of interest to this community as well. Furthermore, that both right occipitotemporal and frontal electrodes also had explained variability that peaked in the sixth CNN layer corroborates the view that a shallower artificial neural network might outperform deeper ones on scene classification tasks.

Although CNNs differ significantly from biological networks, they are of interest to neuroscientists because they allow us to see a solution to a difficult information-processing problem in a step by step manner. The extent to which hard problems such as visual recognition have unique solutions is an open question. Thus, the growing number of similarities between biological and neural networks may indicate that artificial neural networks have honed in on the same solution found by evolution and biology.

## Acknowledgements

This work was funded by National Science Foundation (1736274) to MRG and BCH, and James S McDonnell Foundation (220020439) to BCH.

## SUPPLEMENTARY MATERIAL

Here, we report the results of an internal replication experiment based on the experimental procedure reported in the main article. The experimental procedure was identical to the primary experiment, with the only exception being that the scene stimuli were presented to 500 msec. Likewise, all recording parameters and post-processing routines were identical to those reported in the primary experiment excepting that the EEG waveforms were segmented 100 ms before stimulus onset and 500 ms following stimulus onset (i.e., 600 ms epochs).

**Figure S1:**
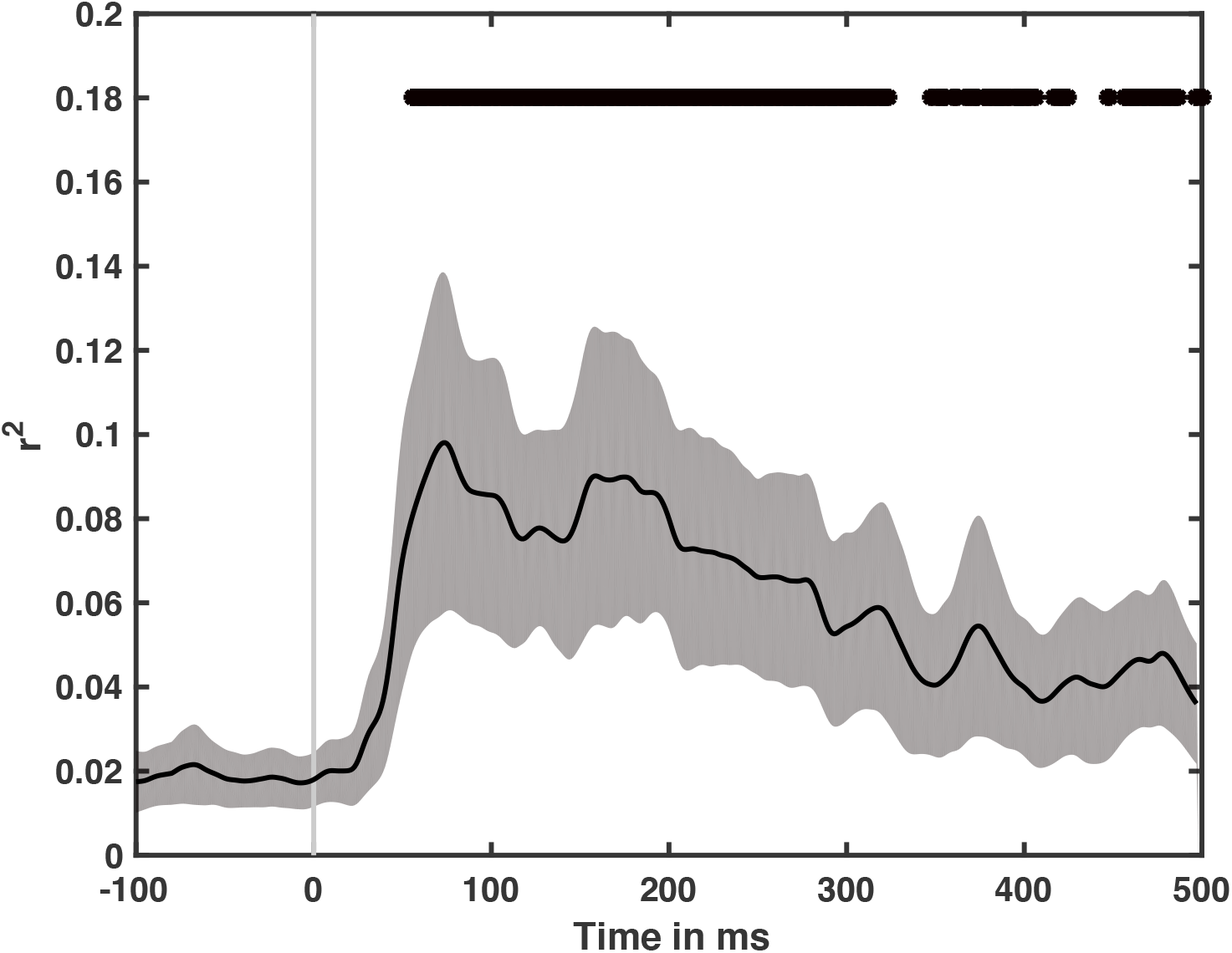
*Overall variance explained by all eight layers of the CNN for the supplementary dataset*.

**Figure S2:**
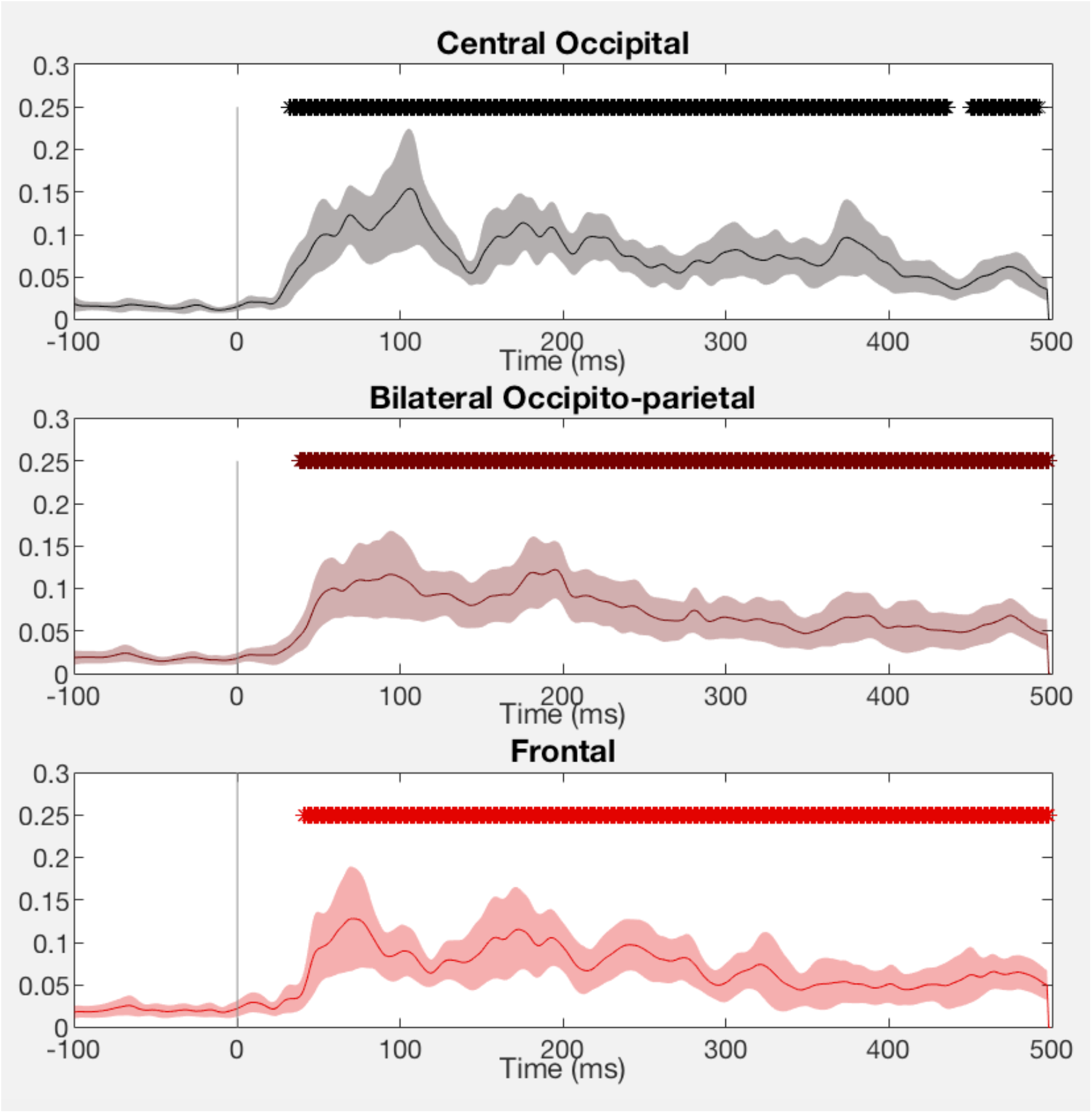
*Overall variance explained by all eight layers of the CNN as a function of electrode group*.

**Figure S3:**
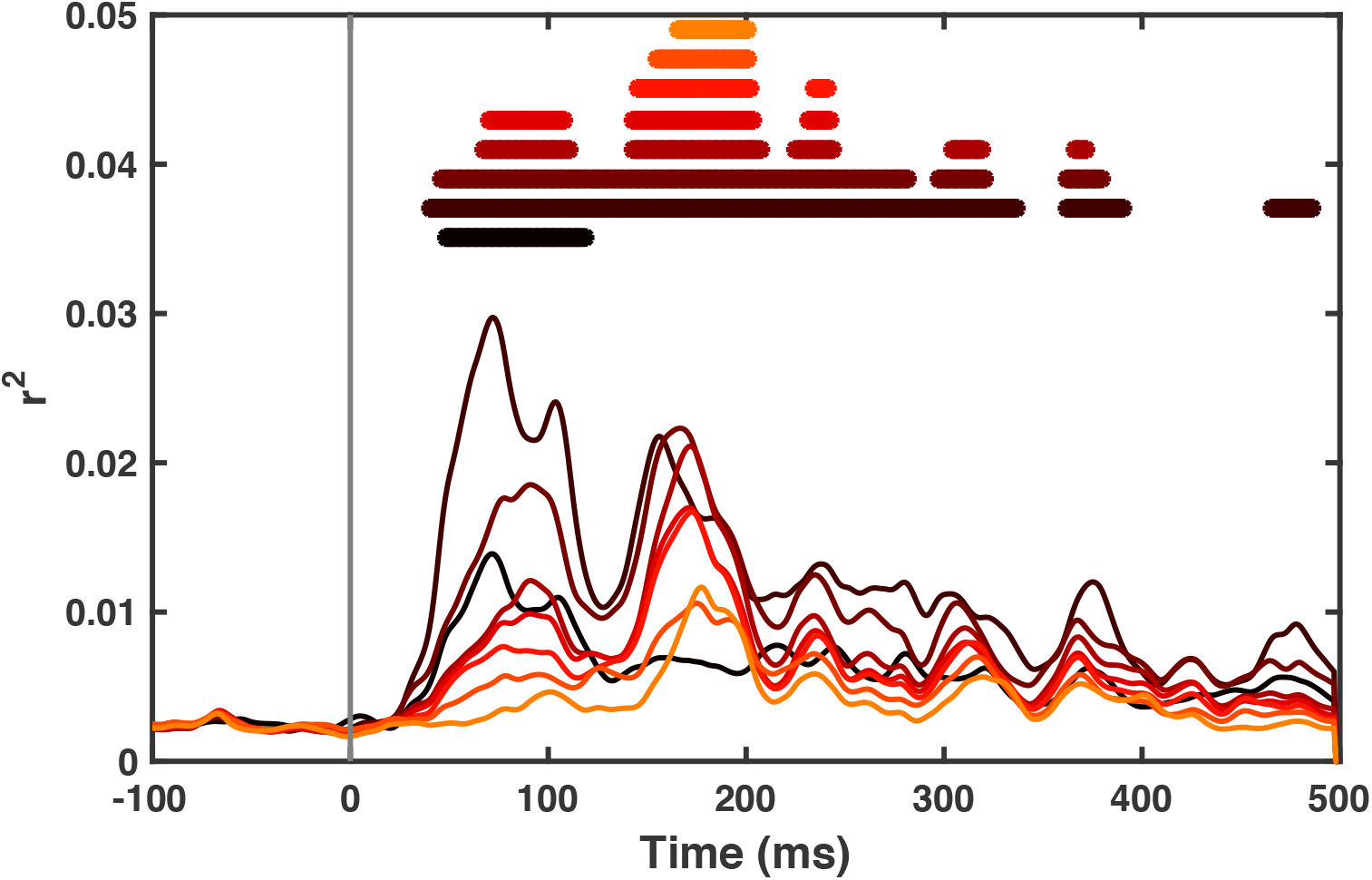
*Variance explained by each layer of the CNN alone*.

**Figure S4:**
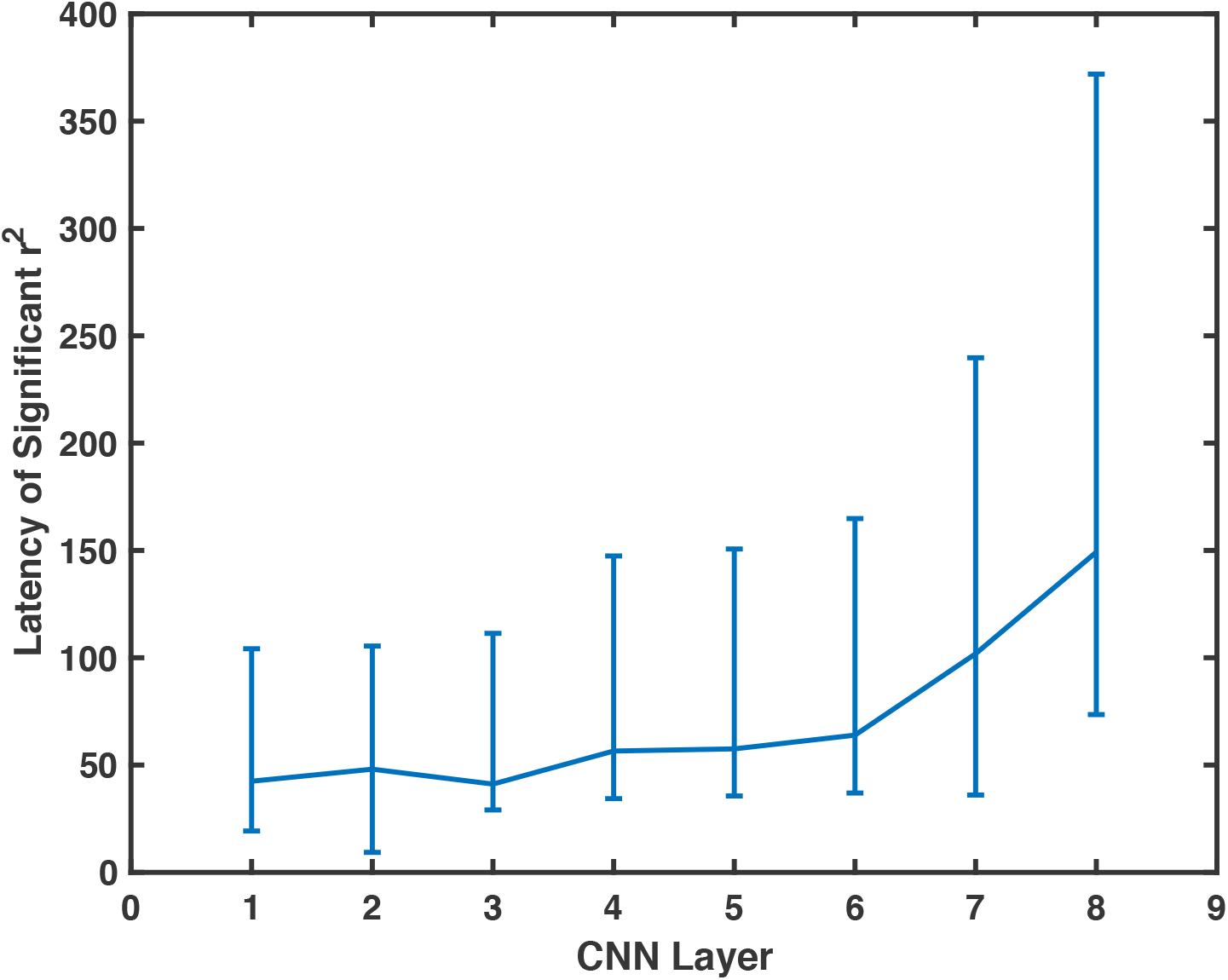
*Onset of significant explained variance as a function of CNN layer*.

## References

Agrawal, P., Stansbury, D., Malik, J., & Gallant, J. L. (2014). Pixels to Voxels: Modeling Visual Representation in the Human Brain. arXiv:1407.5104 [Cs, Q-Bio]. Retrieved from http://arxiv.org/abs/1407.5104

Ba, J., & Caruana, R. (2014). Do Deep Nets Really Need to be Deep? In Z. Ghahramani, M. Welling, C. Cortes, N. D. Lawrence, & K. Q. Weinberger (Eds.), Advances in Neural Information Processing Systems 27 (pp. 2654–2662). Curran Associates, Inc. Retrieved from http://papers.nips.cc/paper/5484-do-deep-nets-really-need-to-be-deep.pdf

Bar, M., Kassam, K. S., Ghuman, A. S., Boshyan, J., Schmid, A. M., Dale, A. M., … Halgren, E. (2006). Top-down facilitation of visual recognition. Proceedings of the National Academy of Sciences of the United States of America, 103(2), 449–454. https://doi.org/10.1073/pnas.0507062103

Bruner, J. S. (1957). On perceptual readiness. Psychological Review, 64(2), 123–152. https://doi.org/10.1037/h0043805

Cadieu, C. F., Hong, H., Yamins, D. L. K, Pinto, N., Ardila, D., Solomon, E. A., … DiCarlo, J. J. (2014). Deep Neural Networks Rival the Representation of Primate IT Cortex for Core Visual Object Recognition. PLOS Computational Biology, 10(12), e1003963. https://doi.org/10.1371/journal.pcbi.1003963

Chang, C.-C., & Lin, C.-J. (2011). LIBSVM: A Library for Support Vector Machines. ACM Trans. Intell. Syst. Technol., 2(3), 27:1–27:27. https://doi.org/10.1145/1961189.1961199

Chauvin, A., Worsley, K. J., Schyns, P. G., Arguin, M., & Gosselin, F. (2005). Accurate statistical tests for smooth classification images. Journal of Vision, 5(9), 659–667. https://doi.org/10.1167/5.9.1

Cichy, R. M., Khosla, A., Pantazis, D., & Oliva, A. (2017). Dynamics of scene representations in the human brain revealed by magnetoencephalography and deep neural networks. NeuroImage, 153, 346–358. https://doi.org/10.1016/j.neuroimage.2016.03.063

Cichy, R. M., Khosla, A., Pantazis, D., Torralba, A., & Oliva, A. (2016). Comparison of deep neural networks to spatio-temporal cortical dynamics of human visual object recognition reveals hierarchical correspondence. Scientific Reports, 6, 27755. https://doi.org/10.1038/srep27755

Cichy, R. M., & Pantazis, D. (2017). Multivariate pattern analysis of MEG and EEG: A comparison of representational structure in time and space. NeuroImage, 158(Supplement C), 441–454. https://doi.org/10.1016/j.neuroimage.2017.07.023

Clarke, A., Devereux, B. J., Randall, B., & Tyler, L. K. (2015). Predicting the Time Course of Individual Objects with MEG. Cerebral Cortex, 25(10), 3602–3612. https://doi.org/10.1093/cercor/bhu203

Delorme, A., & Makeig, S. (2004). EEGLAB: an open source toolbox for analysis of single-trial EEG dynamics including independent component analysis. Journal of Neuroscience Methods, 134(1), 9–21. https://doi.org/10.1016/j.jneumeth.2003.10.009

DiCarlo, J. J., & Cox, D. D. (2007). Untangling invariant object recognition. Trends in Cognitive Sciences, 11(8), 333–341. https://doi.org/10.1016/j.tics.2007.06.010

Edelman, S. (1998). Representation is representation of similarities. The Behavioral and Brain Sciences, 21 (4), 449–467–498.

Fukushima, K. (1988). Neocognitron: A hierarchical neural network capable of visual pattern recognition. Neural Networks, 1(2), 119–130. https://doi.org/10.1016/0893-6080(88)90014-7

Goddard, E., Carlson, T. A., Dermody, N., & Woolgar, A. (2016). Representational dynamics of object recognition: Feedforward and feedback information flows. NeuroImage, 128, 385–397. https://doi.org/10.1016/j.neuroimage.2016.01.006

Greene, M. R., & Fei-Fei, L. (2014). Visual categorization is automatic and obligatory: Evidence from Stroop-like paradigm. Journal of Vision, 14(1), 14–14. https://doi.org/10.1167/14.1.14

Greene, M. R., & Oliva, A. (2009). The Briefest of Glances: The Time Course of Natural Scene Understanding. Psychological Science, 20, 464–472. https://doi.org/10.1111/j.1467-9280.2009.02316.x

Güçlü, U., & Gerven, M. A. J van. (2015). Deep Neural Networks Reveal a Gradient in the Complexity of Neural Representations across the Ventral Stream. The Journal of Neuroscience, 35(27), 10005–10014. https://doi.org/10.1523/JNEUROSCI.5023-14.2015

Hansen, B. C., & Essock, E. A. (2004). A horizontal bias in human visual processing of orientation and its correspondence to the structural components of natural scenes. Journal of Vision, 4(12), 5–5. https://doi.org/10.1167/4.12.5

Hansen, B. C., & Hess, R. F. (2006). Discrimination of amplitude spectrum slope in the fovea and parafovea and the local amplitude distributions of natural scene imagery. Journal of Vision, 6(7), 3–3. https://doi.org/10.1167/6.7.3

He, K., Zhang, X., Ren, S., & Sun, J. (2016). Deep Residual Learning for Image Recognition. In Proceedings of the IEEE Conference on Computer Vision and Pattern Recognition (pp. 770–778). Retrieved from https://www.cv-foundation.org/openaccess/content_cvpr_2016/html/He_Deep_Residual_Learning_CVPR_2016_paper.html

Hochstein, S., & Ahissar, M. (2002). View from the top: Hierarchies and reverse hierarchies in the visual system. Neuron, 36, 791–804.

Jia, Y., Shelhamer, E., Donahue, J., Karayev, S., Long, J., Girshick, R., … Darrell, T. (2014). Caffe: Convolutional Architecture for Fast Feature Embedding. In Proceedings of the 22Nd ACM International Conference on Multimedia (pp. 675–678). New York, NY, USA: ACM. https://doi.org/10.1145/2647868.2654889

Khaligh-Razavi, S.-M., & Kriegeskorte, N. (2014). Deep Supervised, but Not Unsupervised, Models May Explain IT Cortical Representation. PLOS Comput Biol, 10(11), e1003915. https://doi.org/10.1371/journal.pcbi.1003915

Kriegeskorte, N., Mur, M., & Bandettini, P. (2008). Representational Similarity Analysis – Connecting the Branches of Systems Neuroscience. Frontiers in Systems Neuroscience, 2. https://doi.org/10.3389/neuro.06.004.2008

Krizhevsky, A., Sutskever, I., & Hinton, G. E. (2012). ImageNet Classification with Deep Convolutional Neural Networks. In F. Pereira, C. J. C Burges, L. Bottou, & K. Q. Weinberger (Eds.), Advances in Neural Information Processing Systems 25 (pp. 1097–1105). Curran Associates, Inc. Retrieved from http://papers.nips.cc/paper/4824-imagenet-classification-with-deep-convolutional-neural-networks.pdf

Kubilius, J., Bracci, S., & Beeck, H. P. O de. (2016). Deep Neural Networks as a Computational Model for Human Shape Sensitivity. PLOS Comput Biol, 12(4), e1004896. https://doi.org/10.1371/journal.pcbi.1004896

LeCun, Y., Bengio, Y., & Hinton, G. (2015). Deep learning. Nature, 521 (7553), 436–444. https://doi.org/10.1038/nature14539

Nili, H., Wingfield, C., Walther, A., Su, L., Marslen-Wilson, W., & Kriegeskorte, N. (2014). A Toolbox for Representational Similarity Analysis. PLOS Computational Biology, 10(4), e1003553. https://doi.org/10.1371/journal.pcbi.1003553

Potter, M. C., Wyble, B., Hagmann, C. E., & McCourt, E. S. (2014). Detecting meaning in RSVP at 13 ms per picture. Attention, Perception, & Psychophysics, 1–10. https://doi.org/10.3758/s13414-013-0605-z

Riesenhuber, M., & Poggio, T. (1999). Hierarchical models of object recognition in cortex. Nature Neuroscience, 2(11), 1019–1025. https://doi.org/10.1038/14819

Russakovsky, O., Deng, J., Su, H., Krause, J., Satheesh, S., Ma, S., … Fei-Fei, L. (2015). ImageNet Large Scale Visual Recognition Challenge. International Journal of Computer Vision, 1–42. https://doi.org/10.1007/s11263-015-0816-y

Seeliger, K., Fritsche, M., Güçlü, U., Schoenmakers, S., Schoffelen, J.-M., Bosch, S. E., & van Gerven, M. a. J (2017). Convolutional neural network-based encoding and decoding of visual object recognition in space and time. NeuroImage. https://doi.org/10.1016/j.neuroimage.2017.07.018

Xiao, J., Ehinger, K. A., Hays, J., Torralba, A., & Oliva, A. (2014). SUN Database: Exploring a Large Collection of Scene Categories. International Journal of Computer Vision, 1–20. https://doi.org/10.1007/s11263-014-0748-y

Yamins, D. L., Hong, H., Cadieu, C., & DiCarlo, J. J. (2013). Hierarchical Modular Optimization of Convolutional Networks Achieves Representations Similar to Macaque IT and Human Ventral Stream. In C. J. C. Burges, L. Bottou, M. Welling, Z. Ghahramani, & K. Q. Weinberger (Eds.), Advances in Neural Information Processing Systems 26 (pp. 3093–3101). Curran Associates, Inc. Retrieved from http://papers.nips.cc/paper/4991-hierarchical-modular-optimization-of-convolutional-networks-achieves-representations-similar-to-macaque-it-and-human-ventral-stream.pdf

Yamins, D. L. K, & DiCarlo, J. J. (2016). Using goal-driven deep learning models to understand sensory cortex. Nature Neuroscience, 19(3), 356–365. https://doi.org/10.1038/nn.4244

Zhou, B., Lapedriza, A., Khosla, A., Oliva, A., & Torralba, A. (2017). Places: A 10 million Image Database for Scene Recognition. IEEE Transactions on Pattern Analysis and Machine Intelligence, PP (99), 1–1. https://doi.org/10.1109/TPAMI.2017.2723009

Zhou, B., Lapedriza, A., Xiao, J., Torralba, A., & Oliva, A. (2014). Learning Deep Features for Scene Recognition using Places Database. In Z. Ghahramani, M. Welling, C. Cortes, N. D. Lawrence, & K. Q. Weinberger (Eds.), Advances in Neural Information Processing Systems 27 (pp. 487–495). Curran Associates, Inc. Retrieved from http://papers.nips.cc/paper/5349-learning-deep-features-for-scene-recognition-using-places-database.pdf

